# Horizontal mitochondrial transfer from the microenvironment increases glioblastoma tumorigenicity

**DOI:** 10.1101/2021.09.01.458565

**Authors:** Dionysios C. Watson, Defne Bayik, Sarah E. Williford, Adam Lauko, Yadi Zhou, Gauravi Deshpande, Juliana Seder, Jason A. Mears, Feixiong Cheng, Anita B. Hjelmeland, Justin D. Lathia

## Abstract

While dynamic microenvironmental interactions drive tumor growth and therapeutic resistance, the underlying direct cell-cell communication mechanisms remain poorly understood. We identified horizontal mitochondrial transfer as a mechanism that enhances tumorigenesis in glioblastoma. This transfer occurs primarily from brain-resident cells, including astrocytes, and can be appreciated in vitro and in vivo through intercellular connections between GBM cells and non-malignant host cells. The acquisition of astrocyte mitochondria drives an overall enhancement of mitochondrial membrane potential and metabolic capacity, while increasing glioblastoma cell self-renewal and tumor-initiating capacity. Collectively, our findings demonstrate that mitochondrial transfer augments the tumorigenic capacity of glioblastoma cells and reveals a previously unknown cell-cell communication mechanism that drives tumor growth.

**One-Sentence Summary:** Mitochondrial transfer from astrocytes to glioblastoma alters metabolic profile and enhances the tumor-initiation capacity.

## Main Text

Glioblastoma (GBM), the most common primary brain cancer in adults(*1*), is characterized by numerous cell-intrinsic/extrinsic interactions that drive tumorigenesis. In addition to cell-surface and secreted protein/extracellular vesicle interactions(*2, 3*), treatment resistance of GBM is augmented by the formation of cytoplasmic interconnections and junctions among tumor cells(*4*). These cytoplasmic bridges among tumor cells enable exchange of cellular metabolites as well as mitochondria(*4*), which can play a key role in metabolic function and cellular programming in GBM(*5, 6*). However, the downstream impact of mitochondrial transfer on GBM remains poorly characterized. Given the crucial role of microenvironmental interactions in driving GBM pathobiology(*7*), we hypothesized that acquisition of mitochondria from the tumor microenvironment results in metabolic changes and augmented tumorigenicity in GBM cells.

### Tumor cells acquire mitochondria from glial cells

Recent studies established that mitochondrial transfer in mouse models following ischemic stroke was important for neuronal recovery(*8*). To assess whether non-malignant host cells also transfer mitochondria to GBM *in vivo*, we orthotopically implanted GFP-expressing syngeneic mouse GBM models (SB28 and GL261) into transgenic C57BL/6 mice expressing a mitochondria-localized mKate2 fluorophore by fusion to the localization peptide of cytochrome c oxidase 8 (mito::mKate2 mice (*9*)), **Fig. 1A**). Confocal microscopy of GBM tumors from mito::mKate2 mice revealed mKate2^+^ puncta within 15-50% of GFP^+^ GBM cells (**Fig. 1B-C, fig. S1**), demonstrating that host cell mitochondria were being transferred to GBM cells *in vivo*. By co-staining tissue sections with wheat germ agglutinin to highlight glycoprotein-rich structures(*10*), we observed that transferred mitochondria frequently localized at host-tumor interfaces and at the termini of intercellular connections between GFP^+^ GBM cells and GFP^-^ host cells containing mKate2^+^ mitochondria (**Fig. 1B, fig. S2-3**). Thus, cytoplasmic interconnections previously reported to exist among GBM cells(*4*) likely also play a role in organelle transfer between GBM and microenvironment cells.

**Fig. 1.**
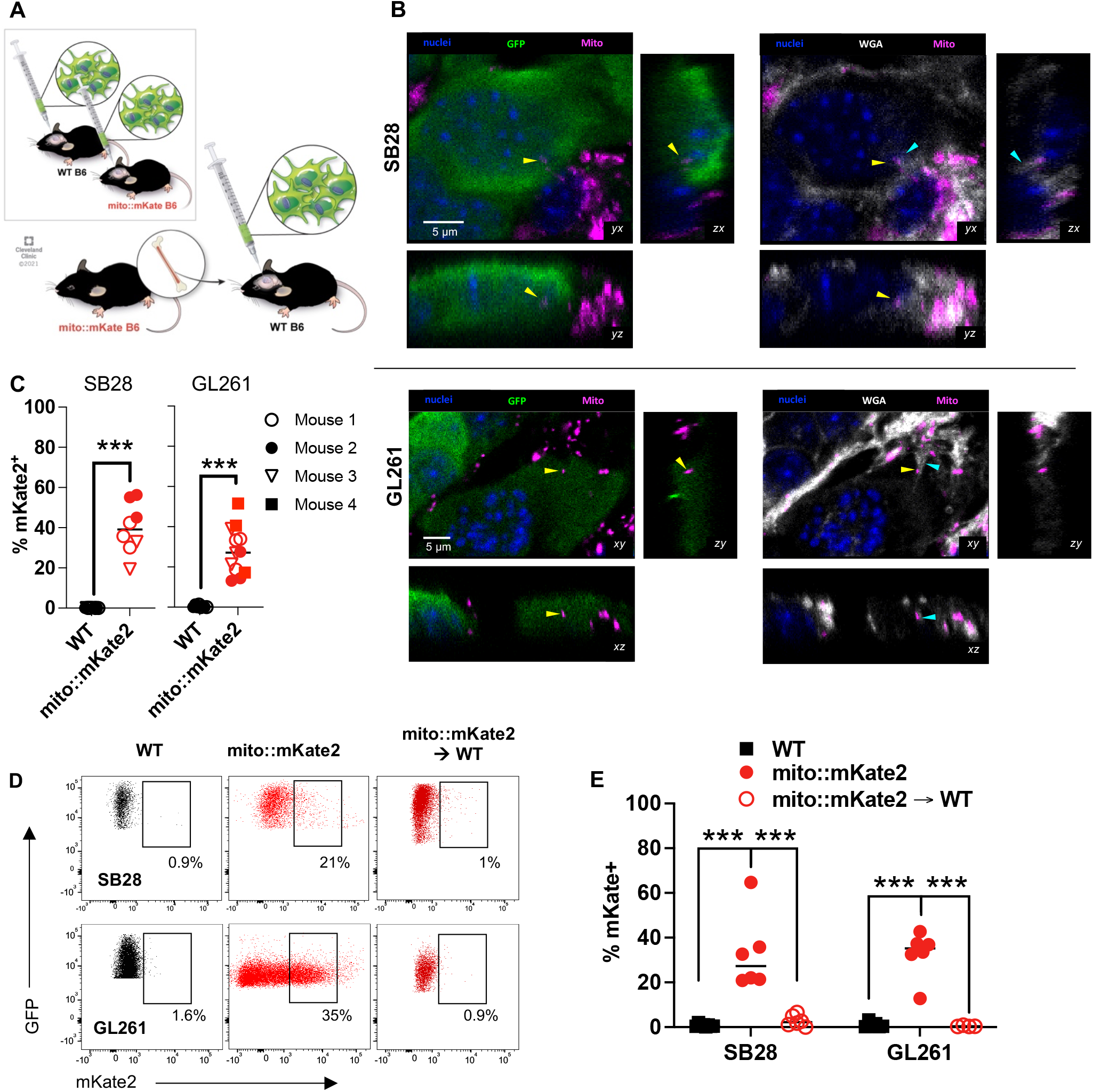
GBM cells obtain host mitochondria from the tumor microenvironment. (**A**) GFP-expressing SB28 and GL261 cells were implanted intracranially into wild-type (WT), mKate2::mito (mKate2) or WT mice with mito::mkate2 bone marrow (mito::mKate2→WT). The assays below were performed at endpoint. (**B**) Single focal planes (xy) and z-stack orthogonal reconstructions (xz, yz) at areas of tumor/host cell interface. Yellow arrowheads point to host mKate+ mitochondria within recipient tumor cells. Cyan arrowheads point to wheat germ agglutinin (WGA)-labelled tether-like structures. (**C**) 3D confocal imaging segmentation-based estimation of mKate2^+^GFP^+^ GBM cell frequency from 2-3 visual fields from n=2-4 mice/group. ^***^ p<0.001, t-test. (**D**) Representative dot plots and (**E**) quantification of relative frequency of mito::mKate2^+^GFP^+^ GBM cells by flow cytometry. ^***^ p<0.001, 2-way ANOVA.

Having observed mitochondrial transfer from the tumor microenvironment to GBM cells in mouse models, we proceeded to interrogate the identity of the host mitochondrial donor cells. GBM tumors in mouse models and patients are known to have significant infiltration by both brain-resident glia and peripheral immune cells that transmigrate into the microenvironment(*7*). We established orthotopic GBM tumors in wildtype C57BL/6 mice that had first received lethal irradiation with subsequent bone marrow reconstitution from mito::mKate2 mice (mito::mKate2→WT) to restrict mKate2 expression to immune cells (**Fig. 1A**). Analysis of single-cell suspensions by flow cytometry indicated negligible host mitochondrial transfer to GFP^+^ GBM cells in mito::mKate2→WT mice, while 20-60% of GFP^+^ GBM cells detected were mKate2^+^ in mito::mKate2 mice (**Fig. 1D-E**). Taken together, these data suggest that brain-resident cells and not tumor-infiltrating immune cells were the major donors of mitochondria to GBM cells *in vivo*.

To further elucidate the identity of the predominant mitochondrial donor cell populations, we co-cultured prevalent tumor-infiltrating cell types with GFP^+^ GBM cells at a 1:2 donor-to-recipient ratio. After 2 hours, we assessed the percentage of mKate2 positivity as a marker of mitochondrial transfer (**Fig. 2A**). Consistent with our *in vivo* observations, we found that brain-resident glia (astrocytes and microglia) donated significantly more mitochondria on a per-cell basis compared to bone marrow-derived macrophages (**Fig. 2B, fig. S4A**). While further polarization of macrophages to an M2- or M1-like phenotype potentially favored increased mitochondrial transfer, the degree of transfer was on average 5-10-fold less than that observed with the brain-resident glia *in vitro* (**fig. S4B**). In addition, culturing GBM cells with donor cell-conditioned medium did not result in mKate2 signal transfer to tumor cells, further suggesting that the mitochondrial transfer is primarily contact dependent (**Fig. 2C, fig. S4C**). Live confocal imaging of astrocyte/SB28 cell co-cultures confirmed the occurrence of mitochondrial transfer at donor-recipient contact sites (**Fig. 2D, Movie S1**), consistent with our *in vivo* observation that host-tumor intercellular connections contained host mitochondria (**fig. S2-3**). Taken together, these results indicate that physical interaction of tumor cells with donor cells is required for effective mitochondrial transfer.

**Fig. 2.**
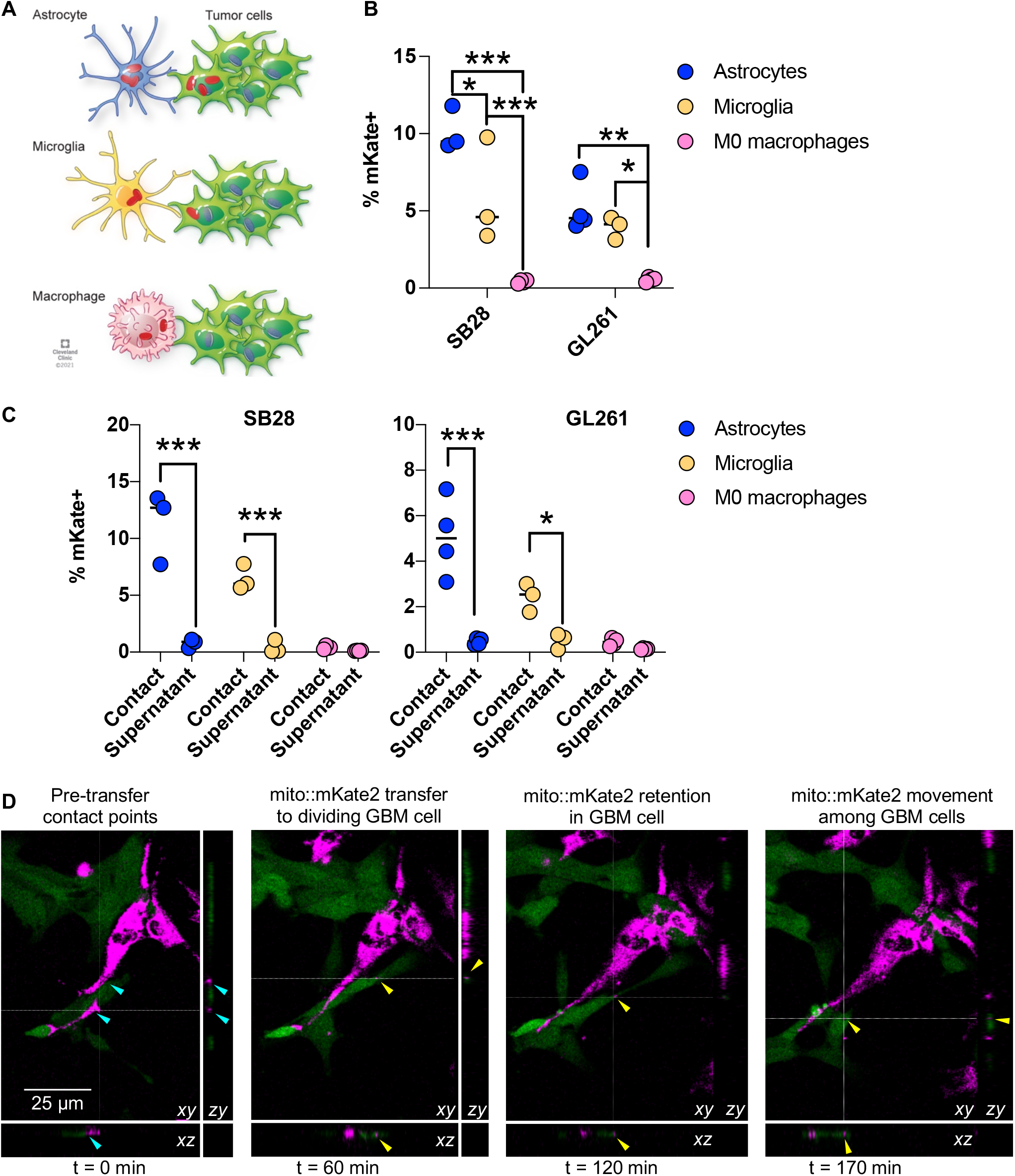
Astrocytes horizontally transfer mitochondria to GBM cells in a contact-dependent manner. (**A**) GBM cells were co-cultured with astrocytes, microglia or macrophages from mito::mKate2 mice for 2 hours, and mitochondrial transfer was analyzed by flow cytometry. (**B**) Relative frequency of mKate2^+^GFP^+^ GBM cells. (**C**) Relative frequency of contact-dependent (contact) versus contact-independent (supernatant) mitochondrial transfer. n=3-4 biological replicates for B, C. ^*^ p<0.05, ^**^ p<0.01, ^***^ p<0.001, 2-way ANOVA. (**D**) Confocal time-lapse images demonstrating real-time acquisition of astrocyte mitochondria (magenta) by SB28 cells (green).

### Mitochondrial transfer impacts tumor cell metabolism

To interrogate the functional significance of mitochondria acquisition, we sorted mKate2^+^ (with exogenous mitochondria) and mKate2^-^ (without exogenous mitochondria) GBM cells from mito::mKate2 astrocyte co-cultures after 2 days (**Fig. 3A**). Unsupervised clustering of RNA-seq analysis revealed that tumor cells had a distinct gene expression profile compared to astrocytes, confirming that sorted mKate2^+^ SB28 cells were not significantly contaminated by astrocytes from the co-culture (**fig. S5A-C**). Additionally, there were a limited number of differentially expressed genes between mKate2^+^ and mKate2^-^ cells, indicating that mitochondrial transfer *in vitro* does not induce broad transcriptional changes (**fig. S5D, Table S1**). However, not surprisingly, genes that were upregulated >1.5-fold in mKate2^+^ samples were enriched within pathways related to mitochondrial biology, in particular electron transport and mitochondrial organization (**Fig. 3B, fig. S6**). In line with this finding, mKate2^+^ GBM cells exhibited increased mitochondrial membrane potential and mass compared to their mKate2^-^ counterparts, as measured by MitoStatus and MitoTracker Deep Red staining (**Fig. 3C, fig. S7A, fig. S7B**). When normalized to total mitochondrial content, mitochondrial potential remained higher in mKate2^+^ cells (**fig. S7A, fig. S7C**).

**Fig. 3.**
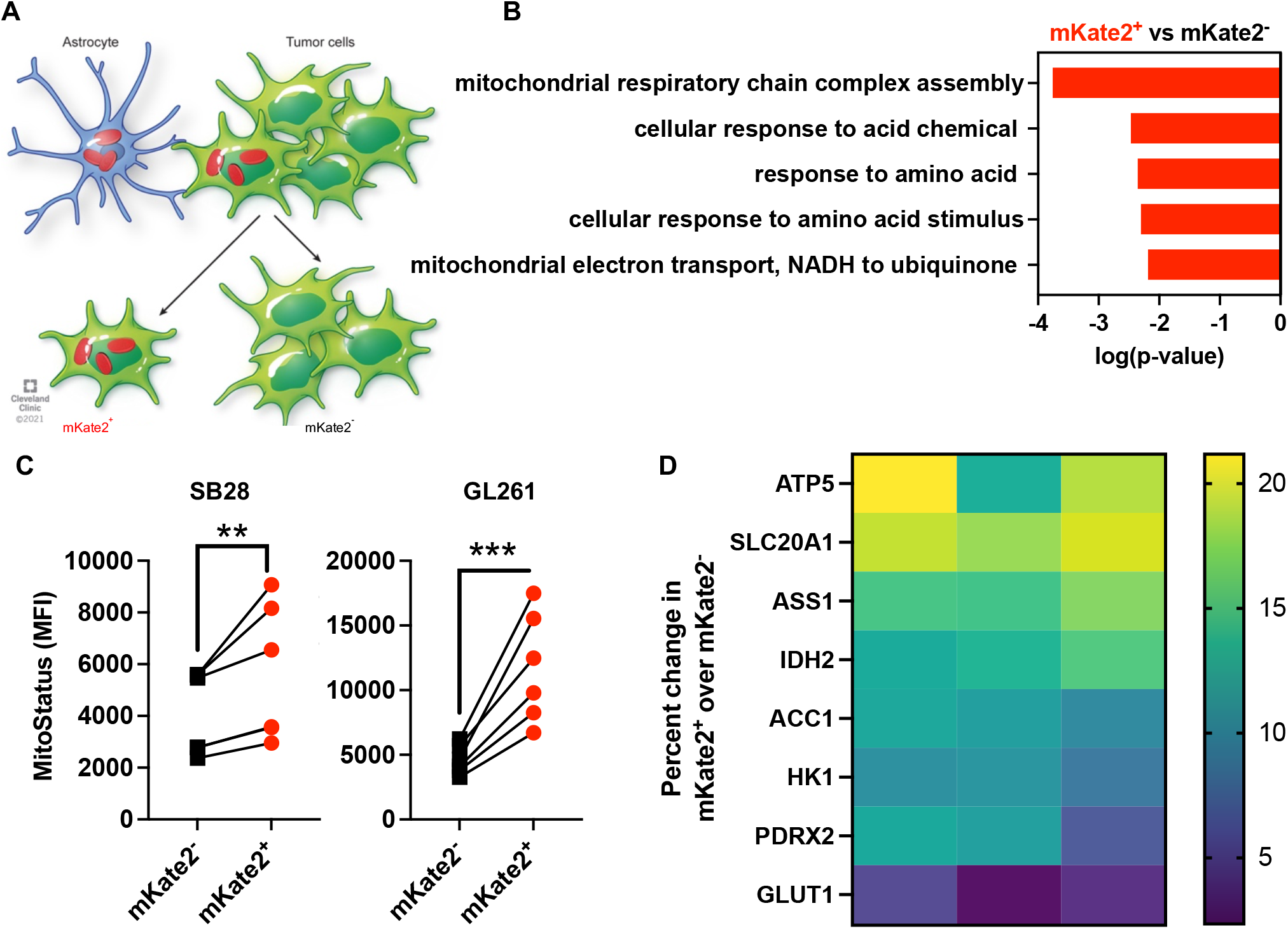
Tumor cells receiving donor mitochondria have altered cellular metabolism. (**A**) GBM cells were co-cultured with astrocytes from mito::mKate2 mice for 48 h. mKate2^+^ and mKate2^-^ SB28 cells were sorted by flow cytometry for downstream assays. (**B**) The top 5 pathways enriched in mKate2^+^ versus mKate2^-^ SB28 cells based on genes upregulated >1.5 fold with p<0.05. (**C**) Geometric mean fluorescence intensity (MFI) of mitochondrial membrane potential dye MitoStatus in mKate2^+^ versus mKate2^-^ GBM cells co-cultured with astrocytes for 48 h. ^**^p<0.01, ^***^ p<0.001, t-test. (**D**) Percent change in MFI of key metabolic pathway proteins in mKate2^+^ versus mKate2^-^ SB28 cells co-cultured with astrocytes for 24 h, as measured by flow cytometry.

To further test the metabolic consequences of acquiring astrocyte mitochondria, we assessed metabolic activity via Seahorse assays, as well as expression of key metabolic proteins by adapting a previously reported metabolic flow cytometry panel(*11*). Metabolic profiling of sorted cells revealed that both mKate2^+^ and mKate2^-^ GBM cells had a lower oxygen consumption-to-extracellular acidification ratio (OCR/ECAR) than matched astrocytes, suggesting that they rely more heavily on glycolytic metabolism in the glucose-rich assay environment, in line with the observed high and invariable GLUT1 expression among these GBM cell subsets (**fig. S8A, 8E, 9**). In contrast, despite similar rates of mitochondrial respiration in the presence of excess glucose, mKate2^+^ cells had higher levels of ATP5A (mitochondrial ATP synthase), SLC20A1 (cellular phosphate import) and ACC1 (fatty acid synthesis), pointing to potential differences in the cellular energetics of mitochondria recipients.

### GBM cells exhibit increased tumorigenicity upon mitochondrial acquisition

To assess whether mitochondrial acquisition impacts the tumorigenic potential of GBM cells, we utilized an *in vitro* limiting dilution assay and found that mKate2^+^ SB28 cells had a higher self-renewal capacity compared to matching cells without exogenous mitochondria, with an estimated stem cell frequency that was ∼2.5-fold higher (**Fig. 4A, fig. S10A**). We further performed an *in vivo* limiting dilution assay by intracranial implantation of decreasing numbers of mKate2^+^ and mKate2^-^ SB28 cells. In three independent experiments, we consistently observed that cohorts receiving mKate2^+^ GBM cells displayed increased penetrance and/or earlier symptomatic disease (**fig. S10B**), corresponding to an estimated 3-fold higher frequency of tumor-initiating cells (**Fig. 4B**). Finally, ∼10-20 % of patient-derived xenograft (PDX) cells also demonstrated the ability to acquire mitochondria from human astrocytes in co-culture, as assessed by flow cytometry and ImageStream in two separate lines (**Fig. 4C, fig. S11**). Taken together, these data demonstrate increased tumorigenicity of cells that received astrocyte-derived mitochondria and suggest that this mode of intercellular communication could play a role in human GBM tumors by altering cellular energetics (**Fig. 4D**).

**Fig. 4.**
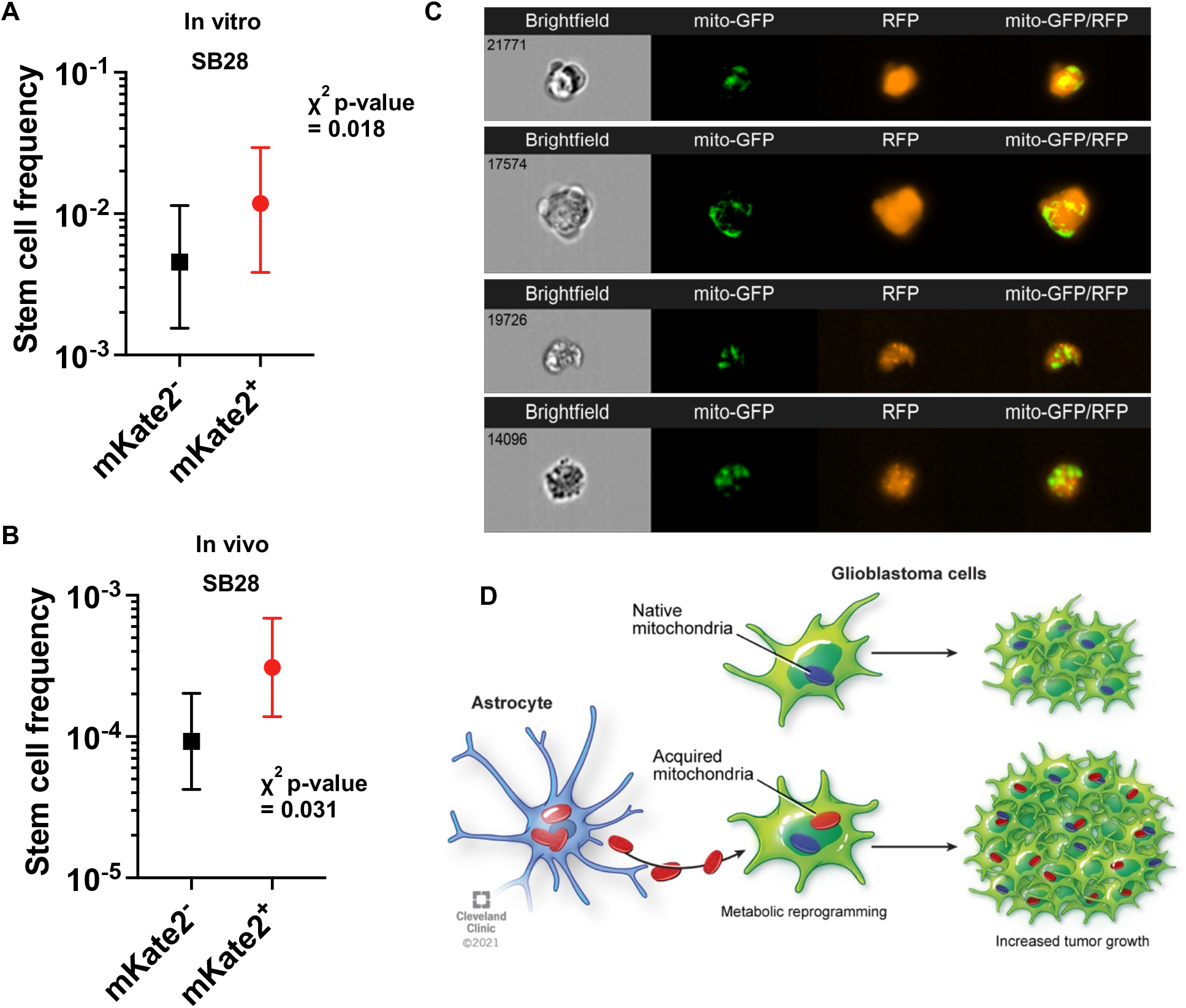
Mitochondrial transfer enhances tumor-initiating capacity. (**A**) Sorted tumor cells were cultured at decreasing cell densities, and the number of wells containing spheres was counted 14 days later. Representative data from n = 3 experiments shown as mean ± 95% confidence interval. (**B**) Decreasing concentrations of mKate2^+^ and mKate2^-^ SB28 cells were implanted into n=4-5 mice. The number of animals that developed tumors within 60 days was used to calculate tumor-forming cell frequency. (**C**) ImageStream was used to visualize transfer of GFP-labelled mitochondria from human astrocytes to RFP-expressing D456 patient-derived xenografts. (**D**) GBM cells acquiring exogenous mitochondria undergo metabolic reprogramming and have enhanced tumorigenicity.

Organelle transfer is an increasingly recognized biological process in models of GBM(*12-14*) and other cancers(*15-17*). However, there is a lack of understanding about its *in vivo* relevance, specific donor-recipient interactions and downstream effects. We found that in the context of GBM, this is a frequent *in vivo* event, involving primarily brain-resident glial cell donors, and is associated with recipient GBM cells acquiring a more tumorigenic phenotype. Mitochondrial transfer led to changes in key metabolic proteins and conferred an advantage to GBM cells within the brain microenvironment. Additional studies into the mechanisms of transfer, as well as into the molecular changes in the recipient cells, are crucial to expand our understanding of this observation and lead to the identification of a new class of therapeutic targets. This line of research could also have broad applicability to tumors outside the central nervous system.

## Acknowledgments

The authors would like to thank Dr. Erin Mulkearns-Hubert for editorial assistance and Ms. Amanda Mendelsohn for illustrations. We would like to thank the Lathia laboratory, LRI Flow Cytometry Core, Imaging Core, Dr. John Peterson, Dr. Anny Mulla and Dr. Prasad Sathyamangla for their technical assistance and discussions.

## Funding

Lerner Research Institute, Case Comprehensive Cancer Center (JDL)

VeloSano Cancer Research Pilot Award (DCW, DB, JDL)

Clinical and Translational Science Collaborative of Cleveland, UL1TR002548 from the National Center for Advancing Translational Sciences (NCATS)

2020 VeloSano Trainee Dream Experiment (DCW)

National Institutes of Health grants 5T32AI007024 and 5TL1TR002549 (DCW)

National Institutes of Health grant K99CA248611 (DB)

National Institutes of Health grant F30CA250254 (AL)

This work utilized the Leica SP8 confocal microscope that was purchased with funding from National Institutes of Health SIG grant 1S10OD019972-01

## Author contributions

Conceptualization: DCW, DB, JDL

Methodology: DCW, DB, SW, AL, YZ

Investigation: DCW, DB, SW, AL, YZ, GD, JS

Visualization: DCW, DB, SW, YZ

Funding acquisition: DCW, DB, JDL

Project administration: JDL

Supervision: JAM, FC, ABH, JDL

Writing – original draft: DCW, DB, JDL

Writing – review & editing: DCW, DB, SW, AL, GD, JS, JAM, YZ, FC, ABH, JDL

## Data and materials availability

Sequencing files have been deposited to GEO with the accession number GSE183004. All other data are available in the main text or the supplementary materials.

## Supplementary Materials

Materials and Methods

Supplementary Text

Figs. S1 to S11

Tables S1

References (18-25)

Movies S1

