## Supplementary Files for "Horizontal mitochondrial transfer from the microenvironment increases glioblastoma tumorigenicity"

**This PDF file includes:**

Materials and Methods
Supplementary Text
Figs. S1 to S11
Tables S1
Captions for Movies S1

**Other Supplementary Materials for this manuscript include the following:**

Movies S1

### Materials and Methods

Mouse tumor cell maintenance and transduction

SB28 cells were gifted by Dr. Hideho Okada (University of California, San Francisco). GL261 cells were obtained from the Developmental Therapeutics Program, National Cancer Institute. All cell lines were treated with 1:100 MycoRemoval Agent (MP Biomedicals) upon thawing and routinely tested for *Mycoplasma spp*. (Lonza). Cells were maintained in RPMI 1640 (Media Preparation Core, Cleveland Clinic) supplemented with 10% FBS (Thermo Fisher Scientific) and 1% penicillin/streptomycin (1% Pen/Strep, Media Preparation Core). For the generation of GFP-expressing GL261 cells, parental GL261 cells were transduced with pReceiver-Lv207 (Genecopoeia) and were selected with 300 µg/ml hygromycin B (Invitrogen). GFP expression was confirmed by flow cytometry.

Mice

All animal experiments were approved by the Institutional Animal Care and Use Committee of Cleveland Clinic and performed in accordance with established guidelines. Tg(CAG-mKate2)1Poche/J (mito::mKate2, stock #032188) mice were purchased from The Jackson Laboratory and were housed in the Cleveland Clinic Biological Research Unit. Both sexes of mito::mKate2 mice were intracranially injected at 4–8 weeks old with 10,000-20,000 SB28 or 100,000 GL261-GFP cells in 5 µl RPMI null media into the left cerebral hemisphere 2 mm caudal to the coronal suture, 3 mm lateral to the sagittal suture at a 90° angle with the murine skull to a depth of 2.5 mm, using a stereotaxis apparatus (Kopf). Age- and sex-matched wildtype littermates were used as controls. Mice were monitored daily for neurological symptoms, lethargy and hunched posture that would qualify as signs of tumor burden.

Bone marrow transplantation

Four-week-old male mice were treated with 11 Gy radiation in two fractions 3-4 hours apart. Reconstitution was achieved by retro-orbital injection of 2 x $10^6$ bone marrow cells from mito::mKate2 mice. Drinking water was supplemented with Sulfatrim (trimethoprim-sulfamethoxazole; Pharmaceutical Associates, Inc.) during the first 10 days, and mice were monitored for an additional 6 weeks for weight loss and symptoms of infection before tumor inoculation. Survival analysis was performed as described above.

Assessment of mitochondrial trafficking *in vivo* by confocal microscopy

At experiment endpoint, mice were perfused with 4% paraformaldehyde in PBS (4% PFA, Fisher Chemical) using the Perfusion One system (Leica). Brains were dissected and stored for 48 h in 20 ml 4% PFA at 4°C, transferred to 20 mL PBS for 48 h at 4°C, and finally transferred to 20 mL 30% sucrose (Fisher Chemical) in PBS at 4°C until density equilibration (typically ~48 h). Samples were embedded in OCT Compound (Tissue-Tek), flash frozen and stored at -80°C. Ten micron sections were prepared using a cryostat (Leica Biosystems), immediately mounted on charged glass slides (Superfrost Plus, Fisherbrand) and stored at -20°C until staining. For staining, selected slides were allowed to warm to room temperature, washed twice with 0.1% Triton-X 100 (Sigma) in PBS (PBS-T) for 10 min, and permeabilized overnight at 4°C with 5% donkey serum (Jackson ImmunoResearch), 0.3% Triton-X 100, 1 mg/mL bovine serum albumin (molecular biology grade, Sigma) in PBS. Sections were then stained with chicken anti-GFP antibody (1:1000 in permeabilization buffer; clone AB_2307313, AvesLabs) overnight at 4°C. Sections were then washed 3 times with PBS-T for 10 min and stained with donkey anti-chicken Alexa fluor (AF)

488-conjugated secondary antibody (1:500 in permeabilization buffer, AB_2340375, Jackson ImmunoResearch) overnight at 4°C. Sections were then washed 3 times with PBS-T for 10 min, then exchanged to HBSS (without phenol red) with $CaCl_2$, $MgCl_2$, $MgSO_4$ and 0.1% Triton-X 100 (HBSS-T) by washing 3 times for 10 min. Sections were immediately stained with wheat germ agglutinin conjugated to AF680 (4 μg/mL in HBSS-T; ThermoFisher) for 1 hour at room temperature and washed 3 times with HBSS-T for 10 min. Nuclear staining was performed with Hoechst 33342 (3.3 μg/mL in PBS-T). Finally, stained sections were mounted and coverslipped with Vectashield Vibrance and no. 1.5 glass coverslips (Fisherbrand). Mounting medium was given at least 24 hours to cure prior to imaging.

Confocal microscopy of stained tissue sections was performed using a Leica SP8. Full-thickness z-stacks were obtained at 3-4 optical fields at sites of tumor-microenvironment interface of each sample using a 63x lens. Still image processing and z-reconstructions were completed using LasX software (version 3.3, Leica). Image analysis for estimation of mito::mKate2 transfer to GBM cells *in vivo* was performed using Velocity software (version 6.3, PerkinElmer). The 3D segmentation algorithms (see **Supplementary Text**) were set to minimize: (a) detection of mKate2-channel noise (using wildtype tissue sections as a negative control) and (b) identification of GFP+ cells not morphologically compatible with tumor cells, likely the result of GFP phagocytosis in the tumor microenvironment (using a relevant size cutoff).

Measurement of mitochondrial trafficking *in vivo* by flow cytometry

At experiment endpoint, mice were euthanized, and resected tumors or the contralateral hemisphere were digested with 1 mg/mL collagenase IV (StemCell Technologies) and 1 mg/mL DNAse I (Roche) for 15 minutes at 37°C. Samples were strained trough a 100 μm strainer (Fisherbrand) and washed with PBS. Cells were stained with LIVE/DEAD™ Fixable Blue Dead Cell Stain Kit (Thermo Fisher Scientific) for 10 minutes on ice, treated with 1:50 diluted FcR blocking reagent (Miltenyi Biotec) for 15 minutes on ice and stained with 1:100 APC-conjugated anti-CD11b antibody (Biolegend, clone M1/70) for 20 minutes to exclude phagocytic cells. Samples were fixed overnight with eBioscience™ FoxP3 Transcription Factor Fixation Kit (Thermo Fisher Scientific) and analyzed with a BD LSRII Fortessa (BD Biosciences) in PBS.

Generation of mouse astrocytes and microglia and *in vitro* mitochondria transfer assay

Brain-resident glial cell cultures were obtained as previously described(*18*). Briefly, mito::mKate2 mice at post-natal day 0-3 were euthanized, and mito::mKate2 signal was confirmed by an EVOS Cell Imaging System (Thermo Fisher Scientific). Brain subventricular zones (SVZs) were digested in 4 mL Accutase Cell Dissociation Reagents (Biolegend) for 5 min and washed with 10 mL neural stem cell (NSC) media: DMEM:F12 (Media Preparation Core), 1% Pen/Strep, 5% FBS, 2% N2 supplement (Thermo Fisher Scientific), 20 ng/mL EGF and 20 ng/mL FGF-2 (R&D Systems). Cells were strained through a 70 μm strainer (Fisherbrand) and cultured into T25 flasks (Corning) in 10 mL NSC media. At confluency, cells were split into two and transferred into T75 flasks (USA Scientific). One flask was kept in NSC media to support astrocyte growth and was split bi-weekly and not used beyond passage 10. The media of the second flask was replaced with microglia polarization media (MPM): DMEM:F12, 1% Pen/Strep, 10% FBS, 20 ng/mL GM-CSF (Biolegend) when the cells reached confluency. Seven days later, microglia growing on the underlying feeder cell layer were shaken off for 1 hour.

A total of 20,000-40,000 astrocytes and microglia were separately cultured in a 96-well flat-bottom plate (Thermo Fisher Scientific) in NSC medium or MPM. Forty-eight hours later, supernatants were collected and centrifuged at 400 *g* for 5 minutes to remove residual cells. Tumor cells were added at a recipient:donor ratio of 2:1. Samples were incubated for 2 hours and treated with Accutase to generate single-cell suspensions. Cells were transferred to 96-well U-bottom plates (Thermo Fisher Scientific) to stain with Live/Dead dye. Exogenous mitochondria uptake of $GFP^+$ tumor cells was assessed with a BD LSRII Fortessa.

<u>Generation of mouse macrophages and *in vitro* mitochondria uptake assay</u>

Bone marrow from the femur and tibia of 4-8-week-old male and female mito::mKate2 mice was flushed with PBS using a 27 G needle. A total of 80,000 cells was cultured in 24-well plates (Corning) and treated with 50 ng/ml recombinant mouse M-CSF (Biolegend) in IMDM (Media Preparation Core) supplemented with 1% Pen/Strep and 20% FBS for 6 days. IFNγ or IL-4 (50 ng/ml; Biolegend) was added for 48 hours to further induce polarization of macrophages to M1- or M2-like macrophages. Supernatants were collected and centrifuged at 400 *g* for 5 minutes to remove residual cells. Tumor cells were added at 2-fold abundance in technical duplicates for a 2-hour incubation. Samples were incubated with Accutase for 5 minutes and transferred into 96-well U-bottom plates for staining with the viability dye and anti-CD11b antibody as described above. Exogenous mitochondria uptake was analyzed from $GFP^+$ tumor cells using a BD LSRII Fortessa.

<u>Confocal microscopy time-lapse imaging</u>

SB28 cells (40,000) were co-cultured with 80,000 mito::mKate2 astrocytes in a glass-bottom 35 mm dish (Mat-tek) overnight. Growth medium was replaced with phenol-red free NSC medium. Multiple full-thickness z-stacks were obtained every 10 min using a Leica SP8 microscope in a 37°C chamber supplemented with 5% $CO_2$, 95% humidity using a 20X/0.8NA objective lens. Time-lapse frames were subsequently analyzed by LasX software.

<u>Sorting of tumor cells from co-cultures</u>

Astrocytes were harvested from flasks by Accutase treatment and stained with a 1:1000 dilution of CellTrace™ Violet Cell Proliferation Dye in PBS at 37°C for 20 minutes. Tumor cells and astrocytes were co-cultured at a 1:1 ratio for 48 hours in NSC media. Samples were sorted into RPMI with 20% FBS and cultured overnight in complete RPMI for subsequent functional assays:

*In vitro limiting-dilution assay*

Sorted tumor cells were cultured at decreasing cell densities (400-25 cells/well of a 96-well plate) over 12 technical replicates in Neurobasal medium: Neurobasal Medium (Thermo Fisher Scientific) with 2% B27 (Thermo Fisher Scientific), 1% Pen/Strep, 1 mM sodium pyruvate (Thermo Fisher Scientific), 2 mM L-glutamine (Thermo Fisher Scientific), 20 ng/mL EGF, and 20 ng/mL FGF-2, and the number of wells containing spheres was counted after 14 days. The online ELDA tool (http://bioinf.wehi.edu.au/software/elda/) was used to calculate stem cell frequency(*19*).

*In vivo limiting-dilution assay*

Sorted $mKate2^+$ and $mKate2^-$ SB28 cells were harvested with Accutase after overnight post-sort culture, pelleted by centrifugation and resuspended in basal RPMI. Cells were counted with trypan blue using a TC-20 cell counter (BioRad) and volume adjusted to achieve decreasing cell concentrations (18,000-3,000) for intracranial implantation. After any volume adjustments, cells were recounted immediately prior to implantation to ensure accurate counts. Subsequently, 4-6-week-old C57BL/6 mice were intracranially implanted with equal numbers of either mKate2$^+$ or mKate2$^-$ SB28 cells as described above. The identity of the implanted cells in mice was then blinded to investigators, who monitored animals daily for neurological symptoms indicative of tumor-related morbidity endpoint.

*Seahorse assay*

A total of 15,000-25,000 astrocytes, mKate2$^+$ and mKate2$^-$ tumor cells were seeded in Seahorse XFe24 Cell Culture Microplates (Agilent) in 250 µL complete RPMI overnight. The media was replaced with phenol red-free RPMI supplemented with 10 mM glucose, 2 mM glutamine and 1 mM pyruvate (Agilent) and incubated for 45 min in a non-CO2 incubator. The oxygen consumption rate (OCR) and extracellular acidification rate (ECAR) were measured using the Seahorse XF24 platform (Agilent).

<u>MitoTracker and MitoStatus staining</u>

CellTrace Violet-stained astrocytes and tumor cells were co-cultured in 6-well plates at a 2:1 ratio for 48 hours. Samples were removed with Accutase. Cells were washed with PBS and stained with MitoStatus Red (BD Biosciences, 1:20,000 dilution) or MitoTracker Deep Red (Thermo Fisher Scientific, 1:100,000 dilution) in complete RPMI 1640 supplemented with 10% FBS and 1% Pen/Strep for 15 minutes at 37°C. Samples were washed with complete RPMI 1640 twice and resuspended in PBS. Sample acquisition was performed using a BD LSRII Fortessa.

<u>Metabolic Flow</u>

The metabolic flow panel was adapted from Ahl et al.(*11*). CellTrace Violet-stained astrocytes were co-cultured with tumor cells overnight at a 1:1 ratio. Samples were fixed in eBioscience™ FoxP3 Transcription Factor Fixation Buffer for 30 minutes on ice and stained with the following antibodies in 1x Permeabilization Buffer for 30 minutes at room temperature: anti-argininosuccinate synthetase 1 (ASS1) [2B10] (Abcam, ab124465, 1:200), anti-ATP synthase F1 subunit alpha (ATP5A) [7H10BD4F9] (Abcam, ab110273, 1:1000), anti-glucose transporter 1 (GLUT1) [EPR3915] (Abcam, ab115730, 1:40), anti-isocitrate dehydrogenase 2 (IDH2) [EPR7577] (Abcam, ab131263, 1:100), anti-glucose 6 phosphate dehydrogenase (G6PD) [EPR20668] (Abcam, ab210702, 1:400), anti-Acetyl-CoA carboxylase (ACC1) [EPR23235-147] (Abcam, ab269273, 1:1000), anti-peroxiredoxin 2 (PRDX2) [EPR5154] (Abcam, ab109367, 1:200), anti-hexokinase 1 (HK1) [EPR10134(B)] (Abcam, ab150423, 1:20), anti-Carnitine palmitoyltransferase I (CPT1A) [EPR21843-71-2F] (Abcam, ab234111, 1:20), anti-SLC20A1 (Thermo Fisher Scientific, 12423-1-AP, 1:200), anti-mouse IgG1, kappa monoclonal [MOPC21] (Abcam, ab18443, 1:200), anti-mouse IgG2b, kappa monoclonal [7E10G10] (Abcam, ab170192, 1:1000) and Rabbit IgG, monoclonal [EPR25A] (Abcam, ab172730, 1:20-1:500). Samples were washed and resuspended in 1x Permeabilization Buffer containing a 1:2000 dilution of Goat Anti-Mouse IgG H&L (Alexa Fluor® 647) (Abcam, ab150119) or Donkey Anti-Rabbit IgG H&L (Alexa Fluor® 647) (Abcam, ab150075). After 30 minutes of incubation at room temperature, cells were washed with permeabilization buffer and resuspended in PBS for analysis with a BD LSRII Fortessa. Geometric mean fluorescence intensity was used to calculate expression levels after subtraction of the background levels from isotype control staining.

Human tumor cell and astrocyte maintenance and culture

Patient-derived xenograft (PDX) D456 was provided by Dr. Darrel Bigner at Duke University, and PDX JX22 was provided by Dr. Jann Sarkaria at the Mayo Clinic. DMEM:F12 containing 10 ng/mL EGF, 10 ng/mL FGF, 1% sodium pyruvate, 2% GEM21 (Gemini Bio), and 1% Pen/Strep was used to culture the PDX lines. A clone of HEK293T cells able to grow in serum-free media (CSC293T) was generated and cultured as previously described(*20*). Normal human astrocytes (NHAs; provided by Dr. Russell Pieper at UCSF) were cultured in DMEM supplemented with 10% FBS, 1% N2 NeuroPlex Supplement (Gemini Bio), 3 ng/mL EGF, and 1% Pen/Strep. CSC293T cells were transfected with psPAX2, pCMV-VSVG, and pLYS1-Mito-GFP or pCMV-RFP using FuGENE HD transfection reagent (Promega). Viral concentration was determined using the Lenti-X qRT-PCR Titration kit (Takara Bio). NHAs were transduced with mito-GFP lentivirus and selected for cells stably expressing mito-GFP using puromycin. D456 and JX22 cells were infected with RFP lentivirus and selected using Blastacidin S (Gibco) to select for stable RFP-expressing cells. Where indicated, cells were sorted for mito-GFP$^+$ or RFP$^+$ with assistance from the Flow Cytometry Core at the University of Alabama at Birmingham.

ImageStream

RFP-expressing D456 or JX22 cells were co-cultured at a 1:1 ratio in the presence of mito-GFP$^+$ NHAs for 24hrs. Samples were imaged at 40x magnification and extended depth of field (EDF). Mito-GFP was acquired on ch02 and RFP on ch04. Ch01 and ch09 are used for brightfield imaging, and ch12 was used for side scatter. A total of 5000 events was recorded, and relevant single color and unstained controls were used. Data was analyzed using IDEAS software (version 6.2; EMD Millipore).

RNA Sequencing

mKate$^+$ and mKate$^-$ SB28 cells and astrocytes from 3 distinct co-cultures (biological replicates) were sorted into multiple 1.5 mL DNA LoBind microtubes (Eppendorf) each containing 700 μL of RLT Plus lysis buffer (Qiagen) supplemented with 1% 2-mercaptoethanol. RNA isolation was performed using the RNEasy Plus Micro kit (Qiagen). Briefly, sorted cell microtubes were brought to room temperature and vortexed vigorously. Cells were further lysed by passing the cell suspension 5 times through a 20 G needle. Downstream isolation was performed per manufacturer's protocol with the following exception. Up to 1400 μL of sorted cell suspension was sequentially loaded onto each DNA binding column. When multiple columns were used for a given sample, elution of DNA was performed sequentially, using 15 μL of RNAse-free water to pool the isolated DNA in a minimal volume.

RNA sequencing and analysis were performed by GENEWIZ (South Plainfield, NJ). Briefly, samples were sequenced using an Illumina HiSeq, 2x150 bp configuration and ≥350 M raw paired-end reads. An average of 41.6 M paired-end reads was sequenced across 9 samples. After Illumina universal adapters were trimmed, the reads were mapped to the *Mus musculus* GRCm38 reference genome using the STAR aligner v.2.5.2b. Unique gene hit counts were calculated by using featureCounts from the Subread package v.1.5.2.

For comparison of tumor cells with astrocytes, genes with an adjusted p-value < 0.05 and absolute log2 fold change > 1 were called as differentially expressed genes using DESeq2. For assessment of differentially up-regulated pathways in mKate$^+$ versus mKate$^-$ SB28 cells, genes that were up-regulated >1.5-fold with a count number of >50 and an unadjusted p-value of <0.05 (**table S1**) were plugged into https://maayanlab.cloud/Enrichr/.

Protein-protein interactions and network visualization

Differential expression analysis was performed using edgeR 3.34(*21*). Genes with count per million greater than 1 in at least 2 samples were used for the analysis. P values < 0.05 were considered significant. The mouse differentially expressed genes (DEGs) were mapped to the human homologs using the NCBI HomoloGene database (https://www.ncbi.nlm.nih.gov/homologene). We then performed the enrichment analysis using Enrichr(*22*) for the entire DEGs and for the up- and down-regulated genes separately.

The protein-protein interactions (PPIs) among the DEGs were extracted using a human protein interactome we built previously(*23*) that contains 17,706 protein nodes and 351,444 PPI edges. We then visualized this protein-protein interaction network using Cytoscape 3.8(*24*). Genes that localize to mitochondria are indicated by diamond node shape based on the Human MitoCarta2.0 database(*25*).

Data representation and analysis

Flow cytometry data were analyzed and generated using FlowJo software (BD Biosciences, v10.7.2). Graphs were generated and statistical analysis were performed using Excel (Microsoft Office, v16.52) or Prism (GraphPad, v9.2.0) software.

**Supplementary Text**

3D segmentation algorithm parameters (GL261)

(1) Define GL261 objects:
  a. Channel 3 (GFP) threshold
    i. intensities: 11-185
    ii. Minimum object size = 10000 cubic microns
  b. Fill holes in objects
  c. Separate touching objects
    i. Object size guide = 1200 cubic microns

(2) Define mitochondria objects
  a. Channel 1 (mito::mKate2) threshold
    i. Intensities 20-255
    ii. Minimum object size = 1 cubic micron
  b. Separate touching objects
    i. Object size guide = 25 cubic microns
  c. Filter population
    i. Volume >= 0.1 cubic microns

(3) Compartmentalize mitochondria objects inside GL261 objects

3D segmentation algorithm parameters (SB28)

(1) Define SB28 objects:
  a. Channel 3 (GFP) threshold
    i. intensities: 20-255
    ii. Minimum object size = 10000 cubic microns
  b. Fill holes in objects
  c. Separate touching objects
    i. Object size guide = 1200 cubic microns
  d. Filter population
    i. Volume > 600 cubic microns

(2) Define mitochondria objects
  a. Channel 1 (mito::mKate2) threshold
    i. Intensities 20-255
    ii. Minimum object size = 1 cubic micron
  b. Separate touching objects
    i. Object size guide = 25 cubic microns
  c. Filter population
    i. Volume >= 0.1 cubic microns

(3) Compartmentalize mitochondria objects inside SB28 objects

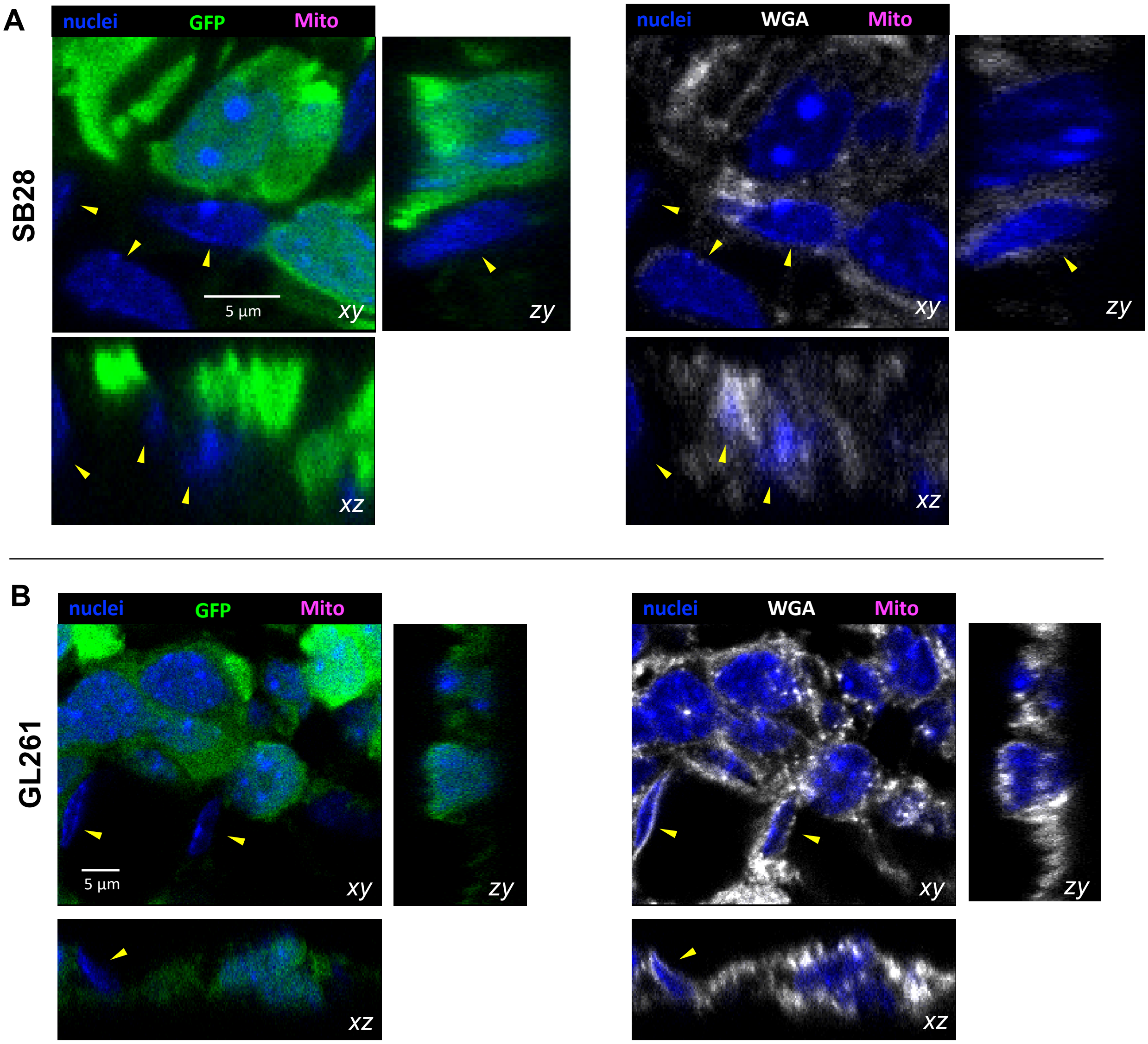


**Fig. S1: Tumor-host cell interface in brain tumor sections of wildtype mice.** GFP-expressing GBM tumor cell lines were implanted intracranially into wildtype C57BL/6 mice. Formalin-fixed, permeabilized cryosections from mouse GBM tumors were stained with anti-GFP, wheat germ agglutinin (WGA) and Hoechst 33342 nuclear stain. Confocal z-stacks were acquired with the detection of the above fluorescent signals, in addition to the mito::mKate2 signal to assess the level of mKate2 background signal at the tumor:host cell interface. Close-up single focal planes and z-stack orthogonal reconstructions of (**A**) SB28 and (**B**) GL261 tumor cells (green) adjacent to host cells (yellow arrowheads).

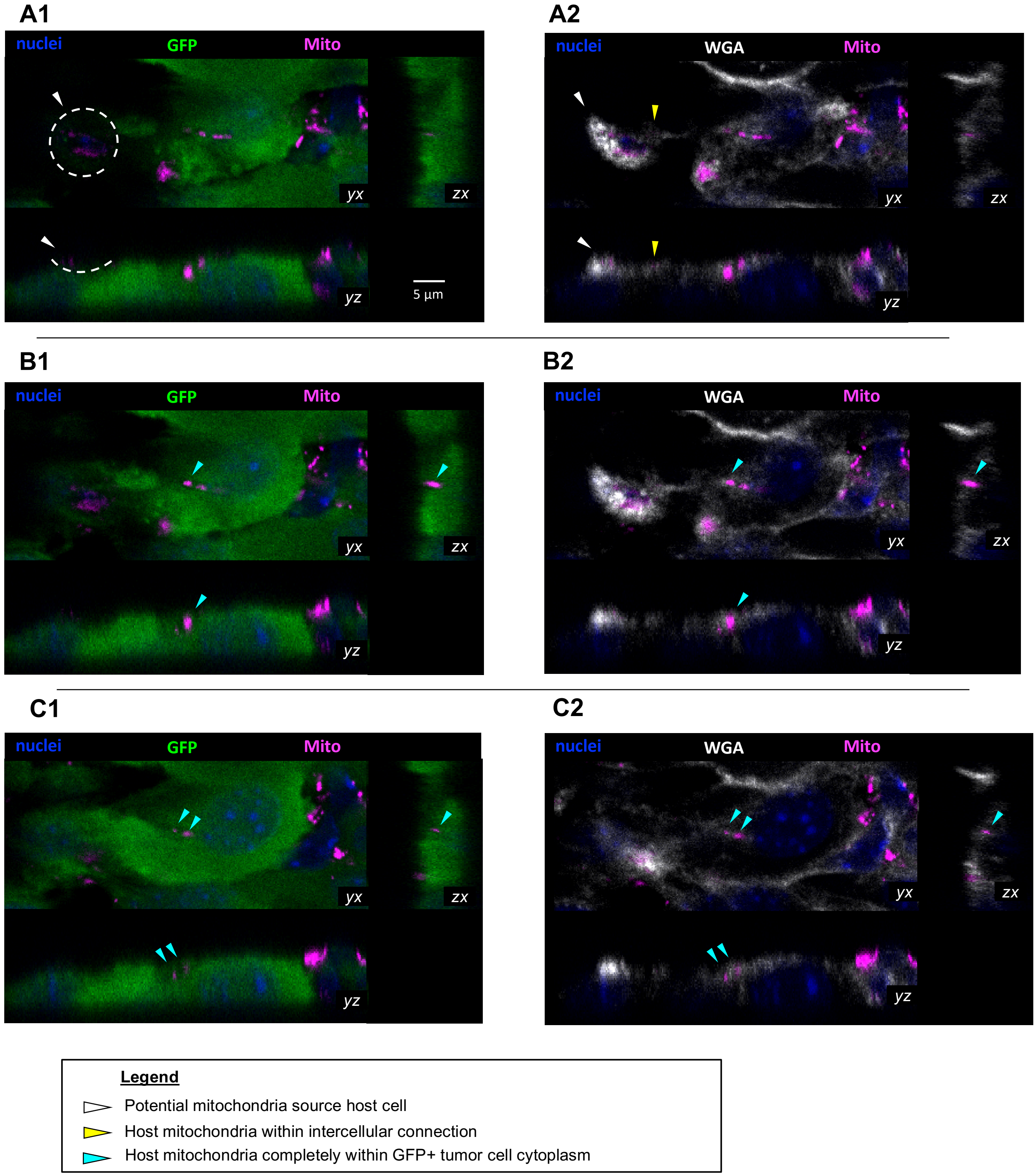


**Fig. S2: Transfer of mitochondria to SB28 cells in vivo.** (**A→C**) Sequential confocal planes (yx) and accompanying orthogonal reconstructions (zx, yz) of SB28 GBM tumors in mice, demonstrating an intercellular connection between a mito::mKate2$^+$ host cell (white arrowheads) and GFP$^+$ tumor cell. Host mKate2$^+$ mitochondria within the intercellular connection are indicated by yellow arrowheads. The mitochondria at the end of the connection are surrounded by GFP signal, suggesting incorporation into the recipient tumor cell cytoplasm (cyan arrowheads).

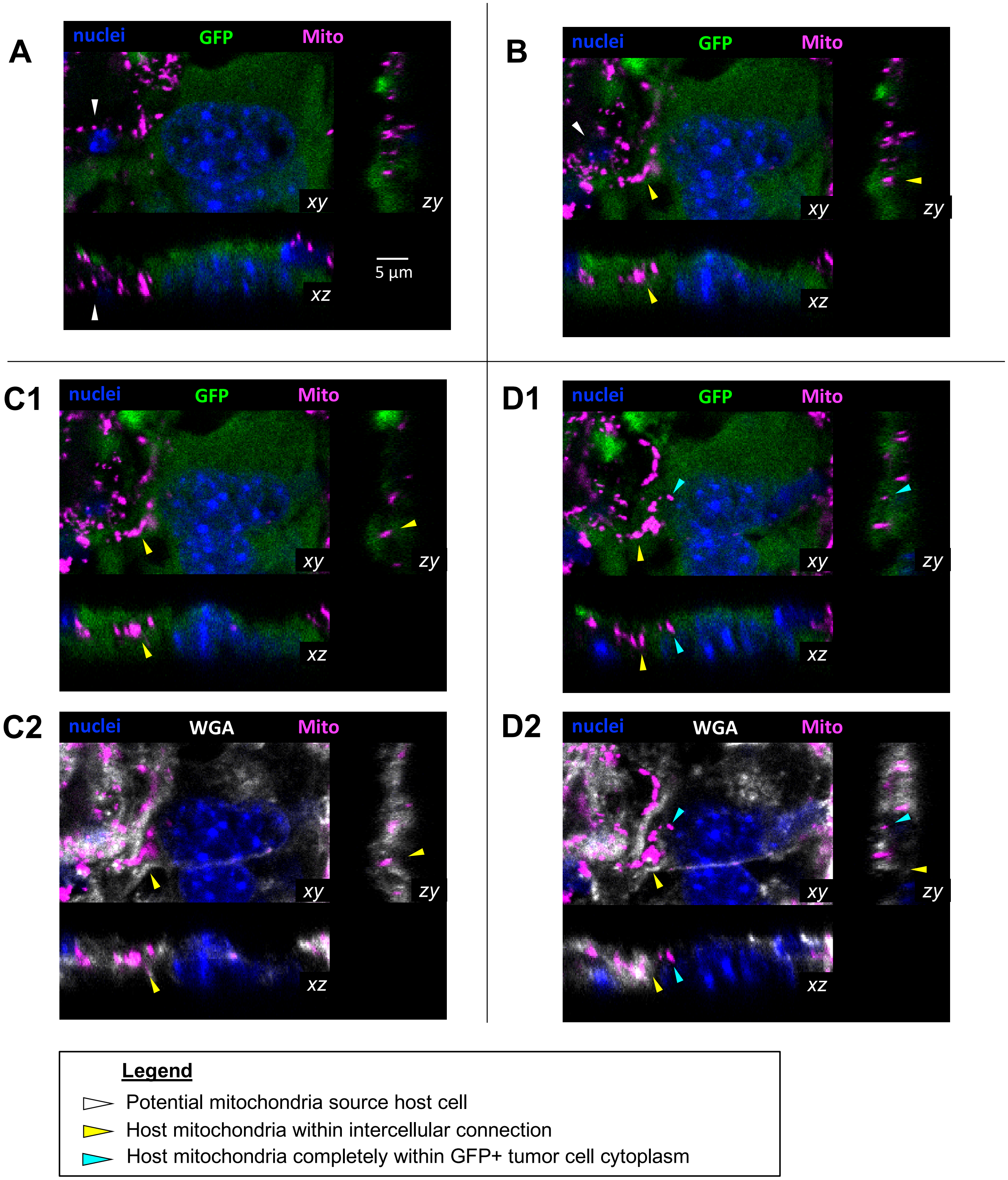


**Fig. S3: Transfer of mitochondria to GL261 cells in vivo.** (**A→D**) Sequential confocal planes (xy) and accompanying orthogonal reconstructions (zy, xz) of GL261 GBM tumors in mice, demonstrating an intercellular connection between a mito::mKate2$^+$ host cell (white arrowheads) and GFP$^+$ tumor cell. Host mKate2$^+$ mitochondria within the intercellular connection are indicated by yellow arrowheads. The mitochondrion at the end of the connection is surrounded by GFP signal, suggesting complete incorporation in the recipient tumor cell cytoplasm (cyan arrowhead).

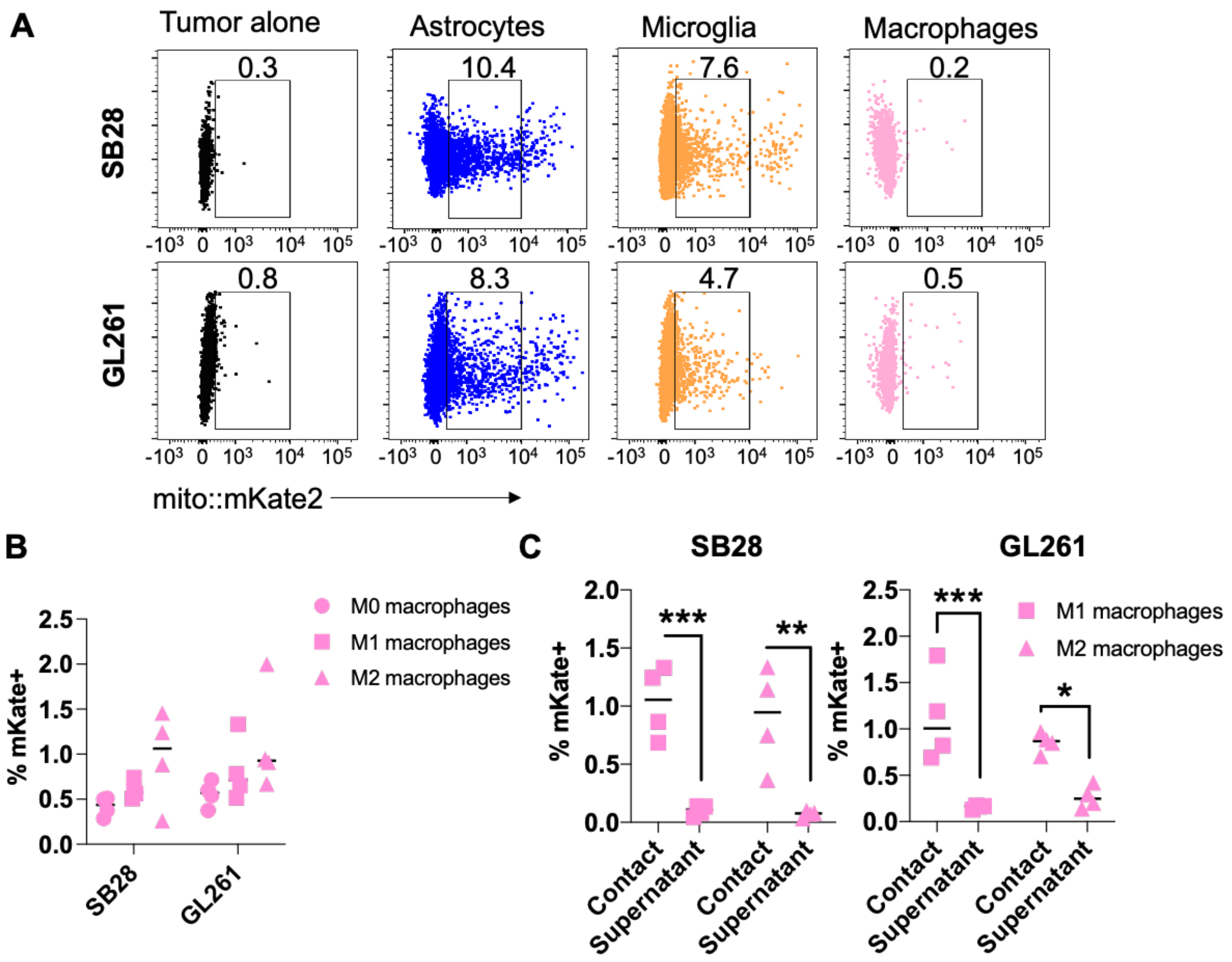


**Fig. S4: Macrophages have a limited contribution to mitochondrial uptake by GBM cells.** (**A**) Representative dot plots depicting the frequency of mitochondrial uptake by GBM cells following 2 hours of co-culture with astrocytes, microglia and macrophages at a ratio of 2:1. (**B-C**) Bone marrow cells were cultured with 50 ng/ml M-CSF for 6 days and treated with 50 ng/ml IFNγ (M1-like) or IL-4 (M2-like) during the last 48 hours. (**B**) mKate2$^{+}$ GBM cell frequency was assessed by flow cytometry after 2 hours of co-culture with the denoted macrophage donors. (**C**) Contact-dependent (contact) and independent (supernatant) mitochondrial transfer was also analyzed by flow cytometry. n=4 biological replicates. * p<0.05, ** p<0.01, *** p<0.001, 2-way ANOVA.

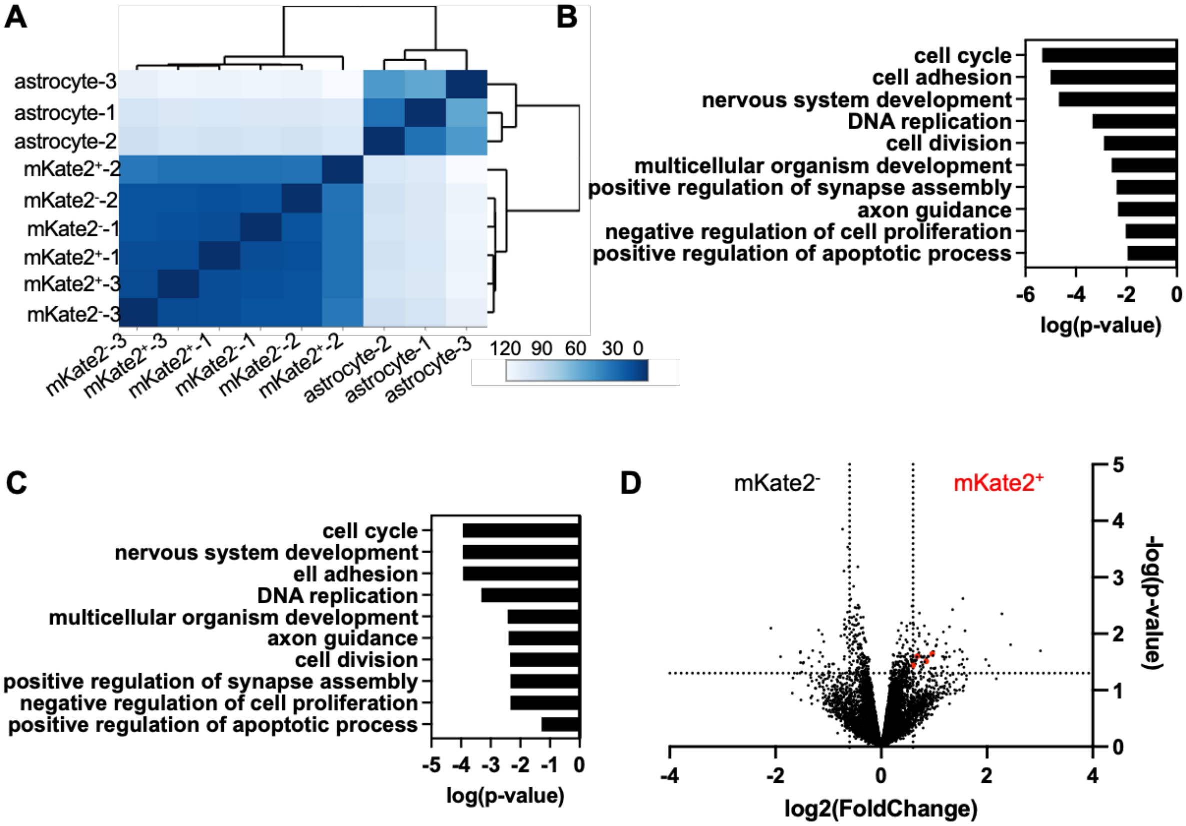


**Fig. S5: Sorted mKate2$^{+}$ cells are not contaminated by astrocytes.** (**A**) Heatmap demonstrating separate unsupervised hierarchical clustering of astrocytes from mKate2$^{-}$ and mKate2$^{+}$ tumor cells based on gene expression profiling. GO pathway analysis of differentially expressed genes between (**B**) mKate2$^{-}$ cells and astrocytes and (**C**) mKate2$^{+}$ cells and astrocytes. (**D**) Volcano plot representing differential gene expression signature of mKate2$^{+}$ versus mKate2$^{-}$ cells. Dashed lines mark fold change > 1.5 and p-value < 0.05. Genes that are mapped to mitochondria-related networks are shown in red.

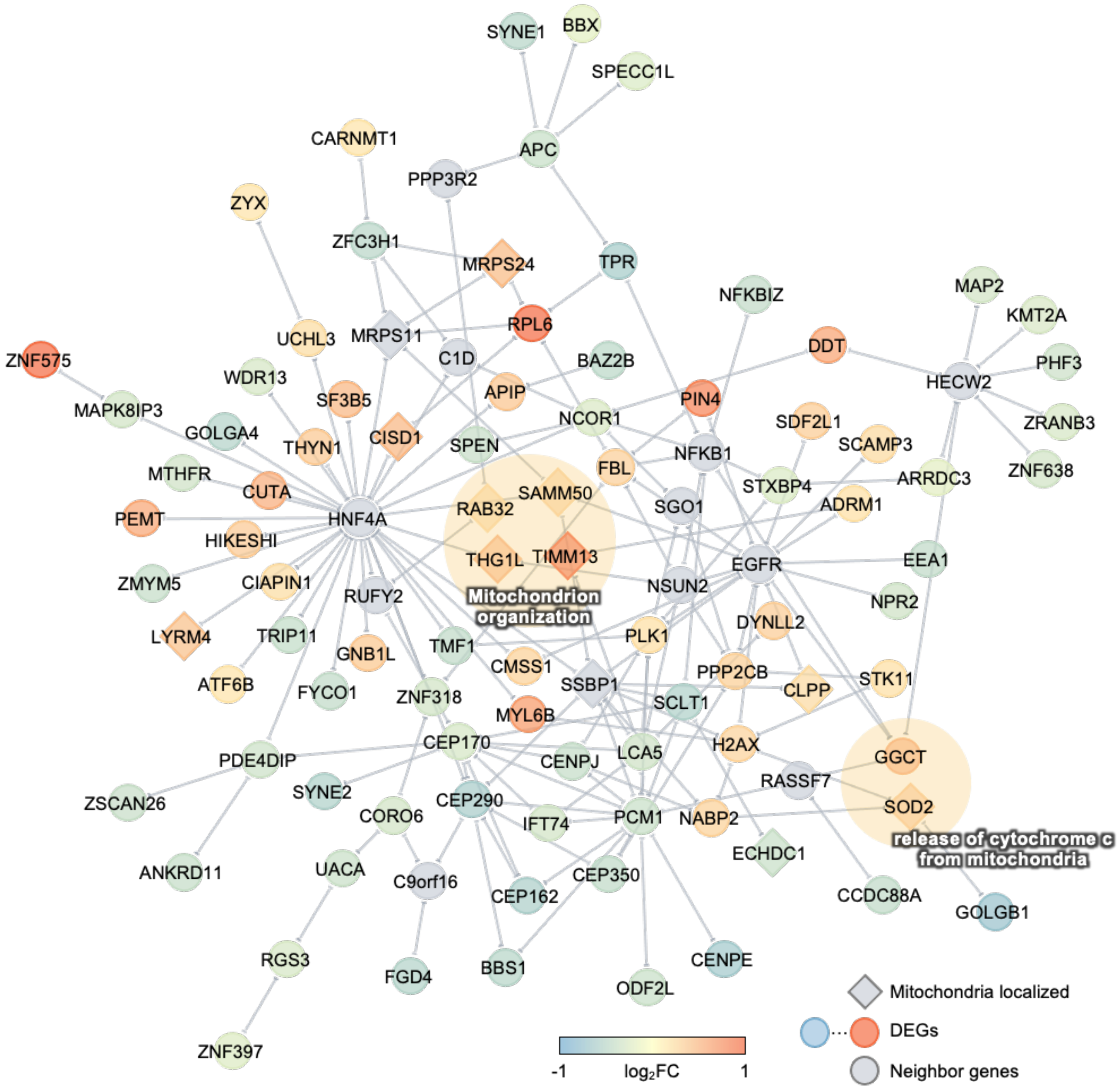


**Fig. S6: Differentially up-regulated genes are enriched in the mitochondrial network.** Protein-protein interaction network for the differentially expressed genes. Mouse genes were mapped to human genes according to NCBI HomoloGene. Protein-protein interactions were extracted for these genes/proteins using our human protein interactome. Diamond-shaped nodes indicate that these genes are mitochondrially localized based on the Human MitoCarta2.0 database. Node color shows the log2 fold change of the genes.

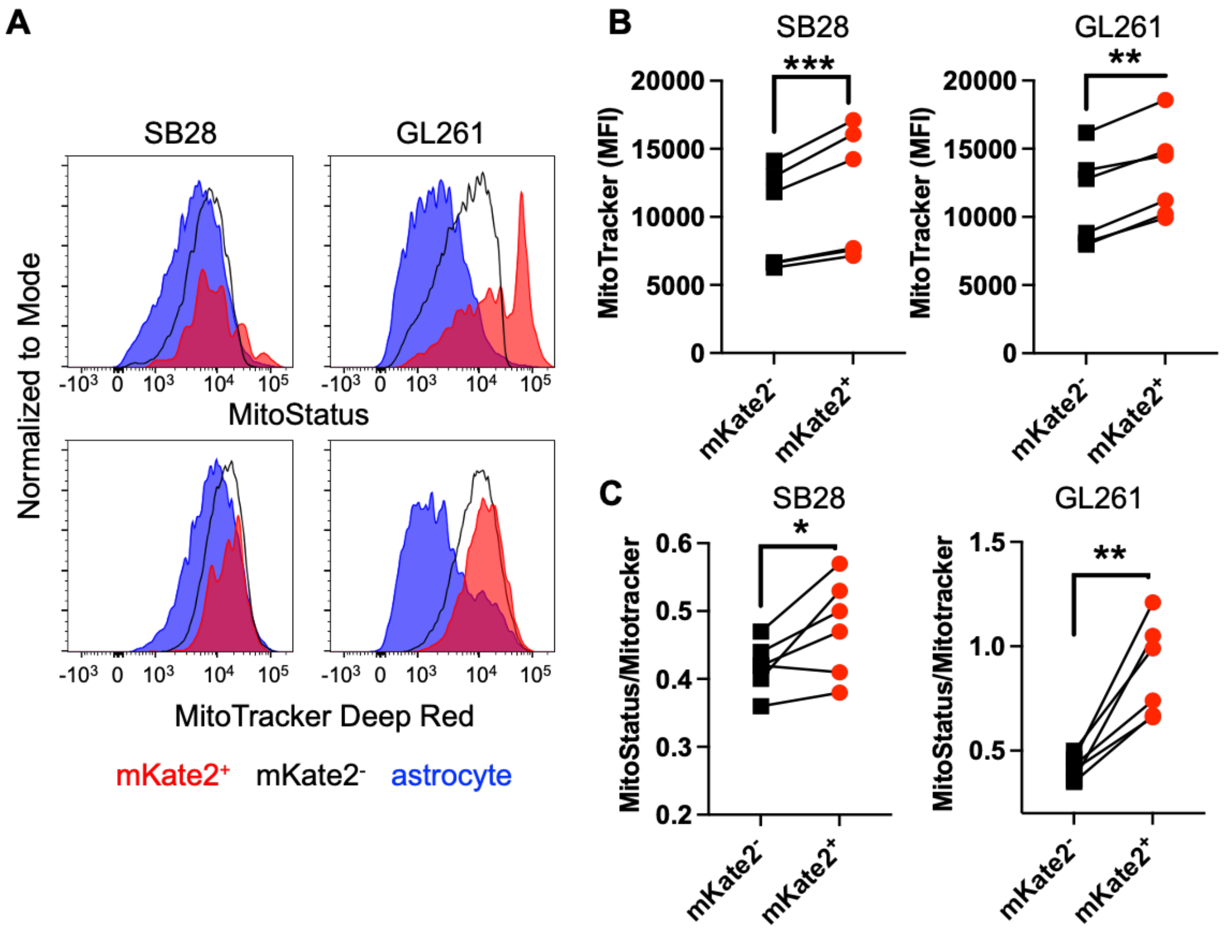


**Fig. S7: Acquisition of donor mitochondria leads to higher mitochondrial content and membrane potential.** (**A**) Representative histograms depicting MitoStatus and Mitotracker Deep Red staining of astrocytes and mKate2$^{+}$ and mKate2$^{-}$ SB28 (left) and GL261 (right). (**B**) Background-corrected Mitotracker Deep Red geometric mean fluorescence of SB28 and GL261 cells revealed an average 15-16% increase in mitochondrial mass in mKate2$^{+}$ compared to mKate2$^{-}$ cells. (**C**) Ratio of MitoStatus to MitoTracker Deep Red staining demonstrated higher membrane potential in mKate2$^{+}$ cells when normalized to total mitochondrial mass. n=6 biological replicates. * $p<0.05$, ** $p<0.01$ and *** $p<0.01$ as determined by paired t-test.

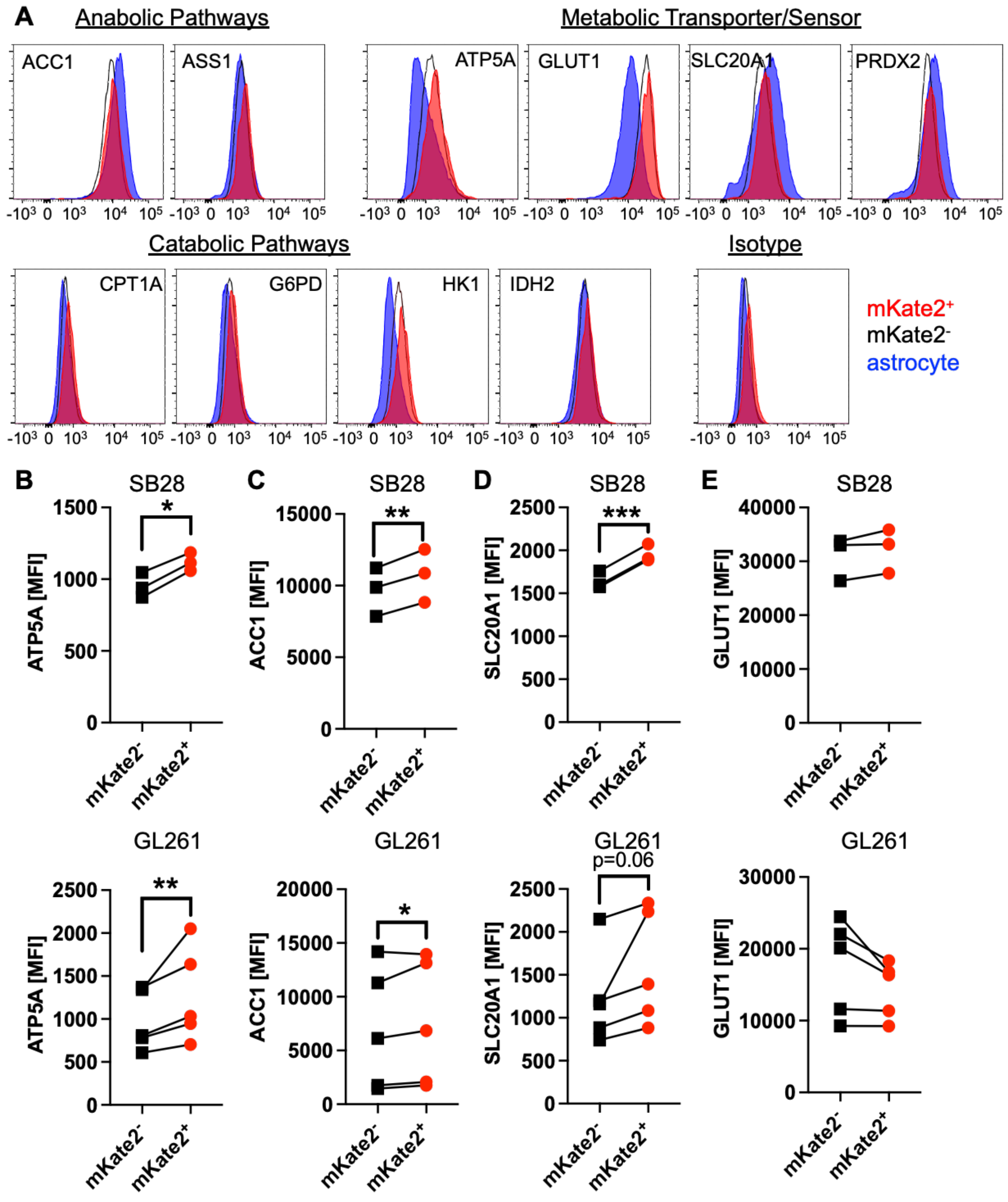


**Fig. S8: GBM cells that acquire astrocyte mitochondria have higher levels of metabolic proteins associated with ATP and fatty acid synthesis.** GBM cell lines were co-cultured with mito::mKate2+ astrocytes for 24 hours and stained with antibodies against key metabolic proteins, as denoted. Expression levels were assessed by flow cytometry. (**A**) Representative histograms depicting differential expression of critical metabolic proteins by astrocytes and mKate2$^+$ and mKate2$^-$ SB28 cells. Astrocytes had higher levels of acetyl-CoA carboxylase (ACC1), SLC20A1 and peroxiredoxin-2 (PRDX2), indicative of more oxidative phosphorylation. In contrast, tumor cells had higher levels of glucose transporter 1 (GLUT 1) and hexokinase 1 (HK1), pointing to a more glycolytic metabolism. (**B-E**) Aggregate data from n=3-5 biological replicates of SB28 (top row) and GL261 (bottom row) cells co-cultured with astrocytes. Panels depict isotype background-corrected, geometric mean fluorescence intensity (MFI) of the expression of the indicated metabolic proteins. * $p<0.05$, ** $p<0.01$, *** $p<0.001$, paired t-test.

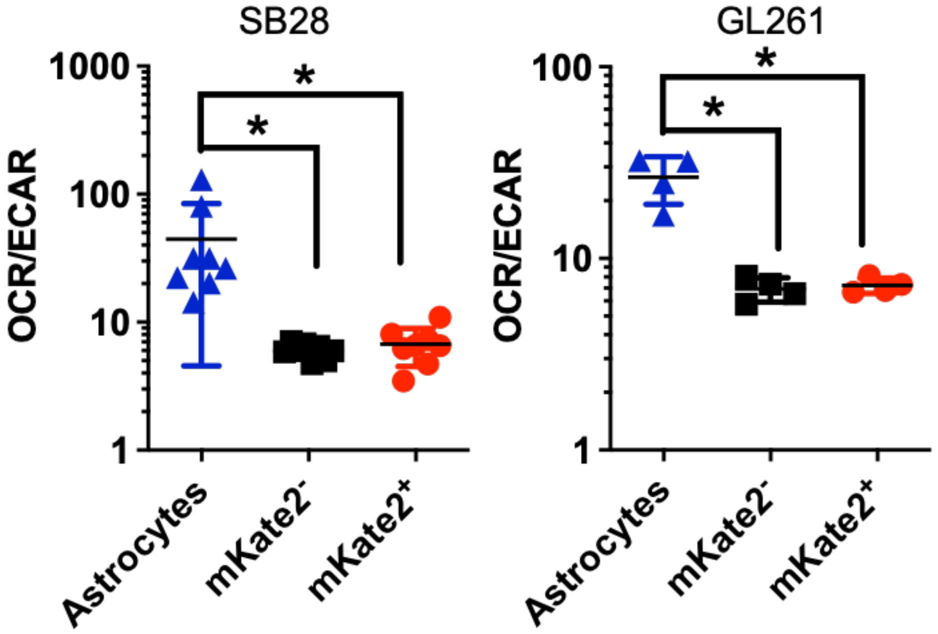


**Fig. S9: mKate2$^+$ and mKate2$^-$ SB28 cells sorted from astrocyte co-cultures similarly rely on glycolysis in a glucose-rich environment.** GBM cells were sorted from mito::mKate2$^+$ astrocyte co-cultures after 48 h, cultured overnight, and then subjected to Seahorse assay in standard assay media in the presence of excess glucose. Oxygen consumption rate (OCR) and extracellular acidification rate (ECAR) were measured at baseline in n=4-8 biological replicates. * $p<0.05$ as determined by one-way ANOVA.

**A**

| | Replicate 1 | | Replicate 2 | | Replicate 3 | |
|---|---|---|---|---|---|---|
| **Group** | **mKate2+** | **mKate2-** | **mKate2+** | **mKate2-** | **mKate2+** | **mKate2-** |
| **1/stem cell frequency** | 39.7 | 45.6 | 84.5 | 220.4 | 68 | 1131 |
| **95% CI** | 26.8-58.8 | 30.9-67.3 | 57.1-125 | 145.7-334 | 44.3-105 | 282.8-4531 |

**B**

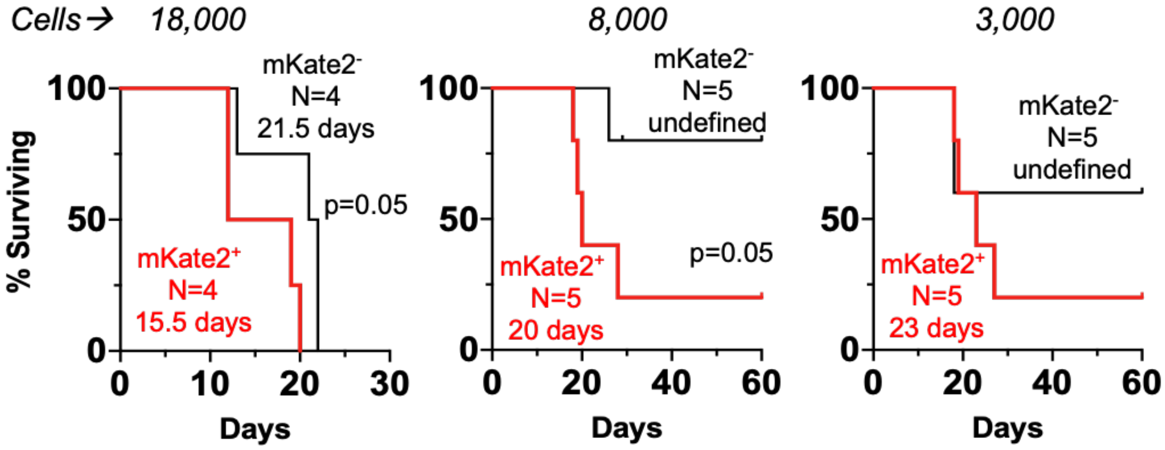


**Fig. S10: mKate2$^+$ SB28 cells have higher self-renewal and tumorigenic capacity.** (**A**) mKate2$^+$ and mKate2$^-$ cells were cultured at decreasing concentrations with 12 technical replicates. Three biological replicates were performed, and stem cell frequency and the confidence interval are shown. (**B**) C57BL/6 mice were implanted with equal numbers of mKate2$^+$ and mKate2$^-$ SB28 cells at decreasing concentrations. Kaplan-Meier curves depict the median survival of tumor-bearing mice, demonstrating that mice implanted with mKate2$^+$ SB28 cells succumbed to tumor earlier. p=0.05 as determined by log-rank test.

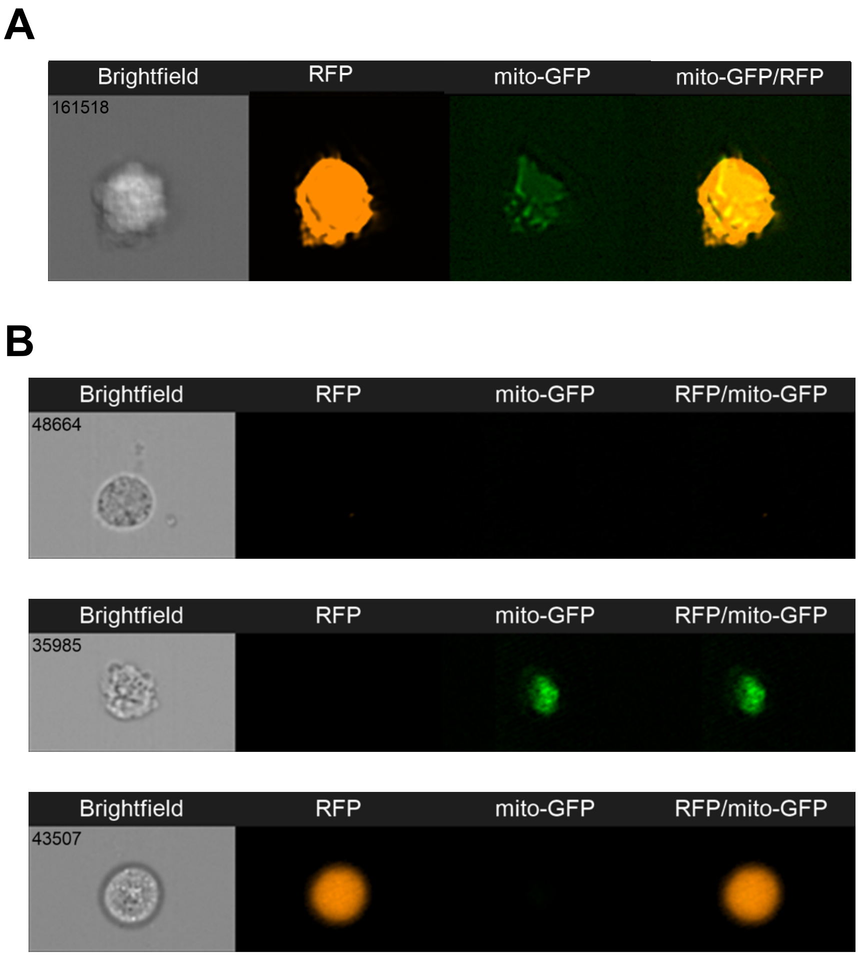


**Fig. S11: PDX cells acquire mitochondria from human astrocytes.** (**A**) PDX line JX22 was incubated with GFP-mito-expressing normal human astrocytes for 24 h at a 1:1 ratio. Mitochondrial puncta in tumor cells were visualized with ImageStream. (**B)** Single-cell controls demonstrating the specificity of the RFP (tumor) and GFP (mitochondria/astrocyte) signals.

| | *Gene List* |
|---|---|
| *Upregulated* | Ndufb11; Ddt; Rnaseh2c; Bola3; Sh3bgrl3; 3110039M20Rik; 1810049J17Rik; Polr2i; Lamtor2; Ndufs5; Myl6b; 2010204K13Rik; Gsta4; Higd2a; 2210008F06Rik; Ropn1l; Timm13; Gm26792; Rpl9-ps6; Pih1d2; Lockd; Lamtor4; Tomm6; Gm9887; G430095P16Rik; Mt2; Snapc5; Snx22; Rpl38-ps2; Pin4; Nrtn; 2010001A14Rik; Immp2l; 2310009A05Rik; Epb41l4aos; Gm11585; Gm13292; Pcbd2; mt-Tn; Smim4; Pet100; Rpl6l; Gm12184; mt-Ta; Gm13436; Serf1; S100a13; Gm8430; Gm2000; Rps26-ps1; Cyp1a1; Gm14586; Gm5905; Gm9843; Tagln; Rps12-ps4; mt-Tc; Gm14303; Gm6204; Mir703; Gm11808; Gm26384 |
| *Downregulated* | Six3os1; Gm21992; Olig2; Rgs8; Zfp874b; Bod1l; Golgb1; Dennd6b; Cntrl; Gm45884; Firre; Mki67; Slc22a17; Zfp940; Gm43737; A930033H14Rik; Gm20721; Akna; Cenpe; Akap9; AC160637.1; Tpr; Cep290; Gm16973 |

**Table S1. Differentially expressed genes between mKate2$^+$ and mKate2$^-$ cells that were used for pathway analysis.** Genes that were up- or down-regulated > 1.5-fold with a p-value < 0.05 and a minimum read count of 50 in mKate2$^+$ SB28 cells compared to mKate2$^-$ SB28 cells.

**Movie S1. Time-lapse confocal images of *in vitro* mitochondrial transfer.** Astrocytes from mito::mKate2 mice were co-cultured with GFP-expressing SB28 cells for 16 hours before live imaging. Full-thickness z-stacks were obtained at 10-minute intervals. Movie depicts a mito::mKate2$^+$ astrocyte (magenta) in contact with a GFP-expressing SB28 cell (green). An mKate2$^+$ mitochondrion is transferred to the dividing GBM cell, which shuttles the mitochondrion back and forth along an intercellular connection linking the GBM daughter cells. At the end of the video, the mKate2$^+$ mitochondrion is retained by the lower daughter cell. See also **Fig. 2D** for still images with accompanying z-reconstructions.
